# Phase-coherence classification: a new wavelet-based method to separate local field potentials into local (in)coherent and volume-conducted components

**DOI:** 10.1101/085605

**Authors:** M. von Papen, H. Dafsari, E. Florin, F. Gerick, L. Timmermann, J. Saur

## Abstract

Local field potentials (LFP) reflect the integrated electrophysiological activity of a large group of neurons. To minimize influence of external activity on the analysis, conventionally bipolar recordings are used to eliminate volume-conducted signals. Here we introduce a novel method, called phase-coherence classification (PCC), to separate LFP in time-frequency domain into a volume-conducted, a local incoherent and local coherent signal. The PCC allows to compute the power spectral densities of each signal and to associate each class with possible locations of electrophysiological activity. In order to test the resolution properties and accuracy of the method we generate composite and non-stationary synthetic time series with similar statistical characteristics as measured LFP. The PCC identifies volume-conducted signals with a phase threshold that is determined from probability density functions of non-phase-shifted synthetic time series. We estimate optimal PCC parameters for the analysis of beta band oscillations in LFP and apply the PCC to a test data set obtained from within the subthalamic nucleus of eight patients with Parkinson’s disease (PD). We show that PCC can identify activity of multiple local clusters during a tremor episode and quantify the relative power of local and volume-conducted signals. We further analyze the electrophysiological response to an apomorphine injection during rest and show that incoherent activity in the low beta band shows a significant medication-induced decrease. We further find significant movement-induced changes on medication of the local coherent signal, which increased during an isometric hold task and decreased during phasic wrist movement. This indicates a different role of incoherent and coherent signals possibly related to physiologically different networks. This new PCC method can potentially also be applied to EEG and MEG data in order to minimize the influence of spatial leakage on power spectra and coherence estimates.

## 1. Introduction

Data obtained from intracranial local field potentials (LFP) using macroelectrodes of ~ 1 mm diameter as well as EEG or MEG data generally reflect the integrated electrophysiological activity of populations of neurons at local and remote locations and mostly stem from post-synaptic potentials (Buzsáki et al., 2012). In order to concentrate on local electrophysiological activity one often uses first or second order derivatives of the measured signal, e.g. bipolar recordings or current source-density for LFP (Mitzdorf, 1985; Lempka and McIntyre, 2013) and average-reference or surface Laplacian for EEG (Hjorth, 1975; Nunez et al., 1997), which reduce the electrodes’ spatial detection range as a large part of the remotely generated and volume-conducted signal is subtracted. Recently, Hipp et al. (2012) introduced a orthogonalization procedure in order to estimate a more localized signal (see also Colclough et al. (2015)). The disadvantage of these techniques is that not only activity generated at distant locations but also highly correlated and non-phase-shifted locally generated activity may be subtracted. Also, the volume-conducted signal is lost for further analyses and incoherent local activity may leak to neighboring electrodes.

The ability to distinguish between local incoherent and coherent signals may help to characterize functionally specialized assembly structures (Schnitzler and Gross, 2005; Denker et al., 2011). Such assembly structures or spatial clusters of focal electrophysiological activity may be encountered in the target structures of clinical applications, e.g. deep brain stimulation of the subthalamic nucleus (STN) in case of Parkinson’s disease (PD). Here focal pathological activity in the beta band (13-30 Hz) (Brown, 2003; Kühn et al., 2004; Hammond et al., 2007; Little et al., 2012) and different topographies of tremor clusters for postural and rest tremor (Reck et al., 2010) were observed in the STN region indicating a functional and patho-anatomical segregation of subloops and symptoms. Further, activity within the beta band (13-30 Hz) may also be subject to physiological modulations, e.g. induced by movement (Engel and Fries, 2010), which need to be differentiated from pathophysiological signals in the same frequency band.

In an attempt to allow for such a differential interpretation of subsignals, namely to identify local (in)coherent and volume-conducted signals in LFP, we developed a novel method called phase-coherence classification (PCC). The PCC separates the signals in time-frequency domain according to their pairwise statistical characteristics into three signal classes associated with electrophysiological activity at local and remote locations. The local signal is further separated into coherent and incoherent activity. With this approach we obtain subsignals, which allow to analyze and estimate the signal components in detail with respect to functionally segregated electrophysiological activity. Conceptually similar wavelet-based separation techniques have been shown to be of great value in the analysis of turbulent flows (Farge et al., 2001; Horbury et al., 2008).

The basic assumption for the application of the PCC is the quasi-static approximation of the electromagnetic field (Plonsey and Heppner, 1967; Stinstra and Peters, 1998). Changes in the extracellular potential propagate across the tissue of the human brain by means of volume-conduction. Therefore, the phase lag of a volume-conducted signal measured at two different locations is negligible. As the sources of LFP and EEG can be approximated as dipoles (Buzsáki et al., 2012; Einevoll et al., 2013), volume-conducted signals generated at remote locations, i.e. postsynaptic terminals, can only generate phase differences of either 0° or 180° between different electrodes. The potential of a dipole source decreases quadratically with distance so that populations close to the electrode have considerably stronger effect on the measurement (Lindén et al., 2011). However, large populations with correlated synaptic input may generate a volume-conducted signal strong enough to be observed at several millimeters distance (Kajikawa and Schroeder, 2011; Lempka and McIntyre, 2013). Note that in our study the interelectrode spacing is 2 mm and, therefore, it is generally unclear how much of the LFP stems from local activity that is only observed at a single electrode and how much of remote activity observed by several electrodes. For the derivation of our method we make use of the quasi-static approximation and assume that signals with phase differences of 0° or 180° between two electrodes are volume-conducted and reflect activity at remote locations (i.e. distances larger than inter-electrode spacings). In contrast signals observed at only one electrode and signals with a phase-shift other than 0° and 180° between electrodes reflect local activity.

In this paper we introduce the wavelet-based PCC and show how to calculate power spectral densities for the separate signals. We focus on the technical aspects of the method, namely its resolution properties and accuracy to determine the correct distribution of power between the subsignals. The resolution of the PCC is tested with synthetic time series and we derive reasonable parameters for an analysis of beta band oscillations in LFP. As a proof of concept and in order to test the accuracy of the method we applied the PCC to non-stationary signals with time varying coherent frequencies. We further analyzed a test data set of LFP measured within the STN of patients with PD. Here, we first show a case study of a tremor episode and then characterize the LFP observed in a cohort of eight PD patients using signals obtained from PCC.

## 2. Materials & Methods

### 2.1. Wavelet Transform

The PCC employs the wavelet transform (Torrence and Compo, 1998) of a discrete time series *x*(*t*_*j*_) with *t*_*j*_ being a discrete point in time defined as

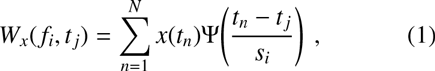

where *N* is the number of data points, Ψ the mother wavelet (here: Morlet) and *s* the temporal scale under consideration. The scale is related to the Fourier frequency *f* according to

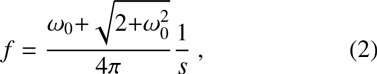

where the wavelet parameter *ω*_0_ defines the number of oscillations in a wavelet and controls the frequency resolution Δ*f*/*f* (Meyers et al., 1993). The frequency resolution can be estimated from the standard deviation of the power spectral density of a monochromatic sine wave.

Using the wavelet cross-spectrum

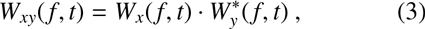

where *W*\* denotes the complex conjugate, the transform allows to compute the instantaneous phase between two signals *x*(*t*) and *y*(*t*) at a given time and frequency according to

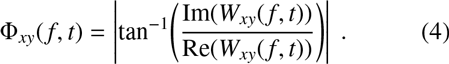

Here Im(*W*) and Re(*W*) are the imaginary and real part of the wavelet coefficient *W*, respectively, and we use the absolute value of the phase because we are only interested in the deviation from 0°.

### 2.2. Wavelet-Based Coherence

The linear correlation between the two signals with respect to magnitude and phase in the time-frequency domain can be quantified by the magnitude squared coherence

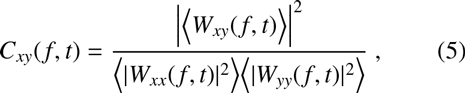

which is the normalized to 0 ≤ *C* ≤ 1. In this study the averaging 〈*〉 was done temporally over a Gaussian with standard deviation

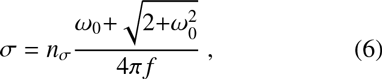

where the averaging width *n*_*σ*_ determines the width of the Gaussian (Grinsted et al., 2004). A value of *n*_*σ*_ = 1 corresponds to the amplitude envelope of the Morlet wavelet. The full width at half maximum of the Gaussian is thus 2.4 ⋅ *n*_*σ*_ *s*.

The significance of the coherence was estimated using a null distribution generated by time series pairs of white noise with unit standard deviation (Lachaux et al., 2002). For comparison we also used pink noise because measured LFP are often observed to have a power law with *f*^−1^. The time series were 26 s long and sampled with Δ*t* = 1/2456 s, which is characteristic of many LFP recordings (Florin et al., 2013). The statistics of the estimated coherence *C*(*f, t*) only depend on the number of non-overlapping segments used for averaging, here determined by the averaging width *n*_*σ*_, and is independent of the length of the time series (Lachaux et al., 2002). We calculated the coherences *C*(*f, t*) between 1000 pairs of noise at 10 Hz and 20 Hz (because we were interested beta band oscillations), for wavelet parameters *ω*_0_ = 6–18 and averaging widths *n*_*σ*_ = 1–10. Subsequently we constructed the probability density functions (p.d.f.) of the null distribution of *C*(*f, t*) from which we determined the 1% significance threshold *C*_1%_ for the coherence.

### 2.3. Wavelet-Based Phase-Coherence Classification

Our novel wavelet-based method, the PCC, is based on the quasi-static approximation (Plonsey and Heppner, 1967; Stinstra and Peters, 1998) and allows to statistically separate a signal *x*(*t*) with respect to a reference signal *y*(*t*) into three different classes. Each wavelet coefficient *W*_*xy*_(*f, t*) is classified as either local incoherent, local coherent or volume-conducted according to the signals’ phase difference Φ_*xy*_ and coherence *C*_*xy*_. For that matter we make the following assumptions:

1) Incoherent activity (red circles in Figure 1): An incoherent signal (*C*_*xy*_ ≤ *C*_1%_) between two electrodes is caused by uncorrelated local activity. It is therefore a marker for the activity in direct proximity of the electrode.

**Figure 1:**
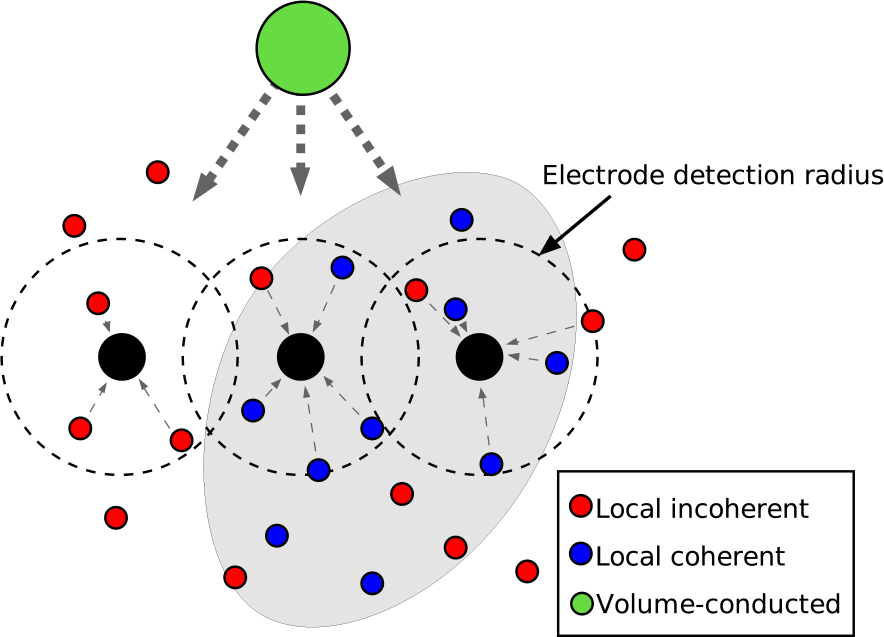
Schematic of three LFP electrodes (black circles) close to a nucleus (gray shaded area) with mostly coherent electrophysiological activity (blue circles). Red circles show locations of incoherent activity and the large green circle denotes the location of volume-conducted activity. Black dashed circles indicate a characteristic electrode detection radius with observed signals denoted by gray dashed arrows. Using PCC we are able to separate the signal power of incoherent, coherent and volume-conducted activity.
2) Coherent activity (blue circles): Coherent signals (*C*_*xy*_ > *C*_1%_) with a non-zero phase difference 0° < Φ_*xy*_ < 180° between two electrodes are caused by correlated local activity. This signal indicates the local activity at multiple, possibly synaptically connected, locations.
3) Volume-conduction (green circle): Signals that are coherent and have a nearly zero phase difference (*C*_*xy*_ > *C*_1%_, Φ_*xy*_ ≈ 0° or Φ_*xy*_ ≈ 180°) are caused by remote activity and are therefore classified as volume-conducted. Note that these signals might also be caused at single local site or two coherent local sites with activity of nearly zero phase difference (type II error).

These classes correspond to certain possible locations of electrophysiological activity as shown schematically in Figure 1. Here, the locations of electrophysiological activity are shown as colored circles and the location of a volume-conducted signal is denoted by a large green circle. The gray shaded area denotes a nucleus that is characterized by mostly coherent activity (as may be the case for the STN in patients with PD). The electrodes’ detection radius is depicted as black dashed circles.

### 2.4. Power Spectral Densities of Signal Classes

The wavelet transform characterizes the spectra of the LFP at a specific time and frequency. Together with PCC we use this localization in time-frequency space in order to separately compute the power spectral densities (PSD) for the local incoherent (*P*_inc_), local coherent (*P*_coh_) and volume-conducted (*P*_vc_) signal. Thus, the LFP can be analyzed in more detail than from mono- or bipolar recordings alone and the changes in the PSD of incoherent, coherent and volume-conducted signals can be further associated with possible locations of electrophysiological activity.

The PSD of the incoherent, coherent and volume-conducted part of a signal *x*(*t*) with respect to a reference signal *y*(*t*) are defined as

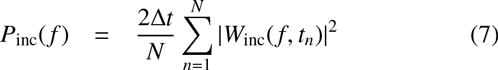

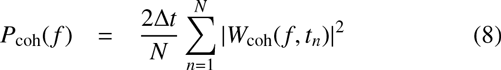

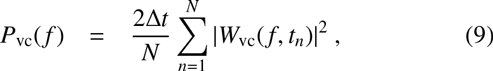

respectively, where the wavelet coefficients *W*_inc_, *W*_coh_, and *W*_vc_ are determined according to the following criteria

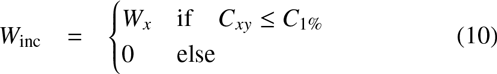

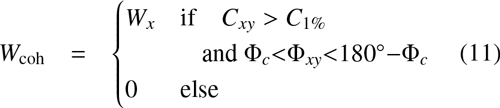

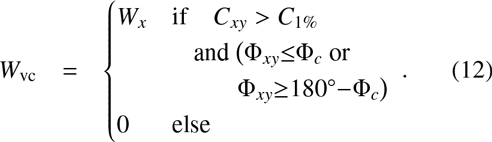

Here Φ_*c*_ is the phase threshold used to define volume-conducted signals (see Section 3.3). Note that in order to relate coherences *C*_*xy*_(*f, t*) and phase differences Φ_*xy*_(*f, t*) we applied the same temporal averaging to the phases as for the coherence estimate.

In order to estimate the average power at a certain electrode independent of the reference electrode one may average the PSD according to

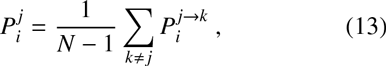

where *N* is the number of neighboring electrodes and 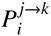 denotes the power of signal class *i* at electrode *j* with respect to reference electrode *k*. From expressions (7)-(9) it follows that the total PSD of signal *x*(*t*)

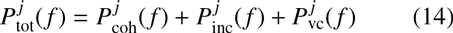

could be obtained from the sum of the separate PSDs of the respective signal classes.

### 2.5. Phase Threshold of PCC

The resolution of the phase difference Φ_*xy*_ and coherence *C* is controlled by the parameters *ω*_0_ and *n*_*σ*_. The larger *ω*_0_ and *n*_*σ*_ the more accurate is the estimation. This, however, comes at the expense of time-resolution (cf. Eq. (6)) so that our aim was to find a reasonable trade-off for these parameters. In order to quantify the phase resolution properties of the wavelet transform we computed the phase differences Φ_12_ between 1000 pairs (*i* = 1, 2) of non-phase-shifted synthetic time series

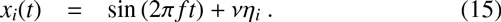

For the determination of the phase threshold in Section 3.3 we used a sinusoidal signal of *f* = 20 Hz, white/pink noise background *η*_*i*_ with unit standard deviation (SD) and a noise level of *v*. The length of the synthetic time series was *T* = 26 s and the sampling rate Δ*t* = 1/2500 s. We used pink noise because the PSD of the observed LFP and many other electrophysiological measurements show an approximate 1 / *f* scaling (He, 2014).

After wavelet transformation of the signals *x*_*i*_(*t*) we calculated the pairwise phase difference Φ_*xy*_ at frequency *f* in order to estimate the phase resolution of our method. In order to relate the phases to the estimated coherence we used the same temporal average controlled by *n*_*σ*_. We computed the p.d.f. of these phase differences Φ_*xy*_ as a function of the parameters *ω*_0_ and *n*_*σ*_. Due to the added noise these were generally nonzero. We then defined the phase threshold

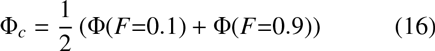

as the mean phase, where the cumulative distribution function

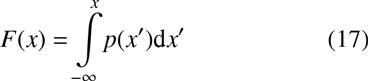

reached *F* = 0.1 and *F* = 0.9. Thus, the threshold correctly classified 80% of the non-phase-shifted signals as volume conducted. Note that the phase threshold is a trade-off between correctly classified volume-conducted signals and coherent signals with small phase-shift. Therefore, phase thresholds defined at higher p-values, e.g. *F* = 0.025/0.975 corresponding to a two-tailed 5% threshold, are not generally a better choice.

### 2.6. Composite Synthetic Time Series

In order to test the accuracy and performance of the PCC we used pairs of synthetic time series with prescribed phase and coherence relations. For that matter we simulated a composition of sinusoidal signals added to a background of pink noise according to

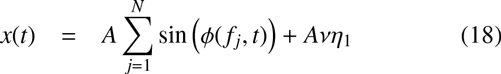

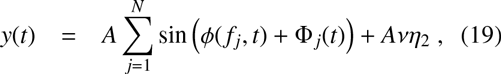

where *N* is the number of sinusoidal signals and Φ_*j*_(*t*) the (time dependent) phase difference between the signal pairs. The phase associated with time dependent frequency *f*_*j*_ is determined according to

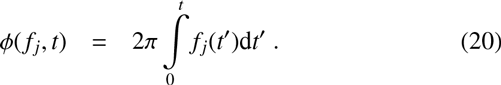

Note that for constant frequency *f*_*j*_ this reduces to 2*πf*_*j*_*t*.

For the accuracy test of the PCC presented in Section 3.5 we used three signals: one centered at *f*_1_ = 10 Hz and only active during 4 s ≤ *t* ≤ 16 s, a second with drifting frequency *f*_2_(*t*) = 20 Hz+*t*/*T* ⋅ 10 Hz, and a third centered at *f*_3_ = 50 Hz. The corresponding phase differences were Φ_1_(*t*) = 30° during 4 s ≤ *t* ≤ 16 s, Φ_2_(*t*) = 0° for *t* ≤ 10 s and Φ_2_(*t*) = 30° for *t* > 10 s, and Φ_3_(*t*) = 30° for *t* ≤ 10 s and Φ_3_(*t*) = 0° for *t* > 10 s. The choice for these parameters reflected the expectation that electrophysiological recordings are non-stationary and therefore some signals may emerge only part of the time while other signals may change their frequency and/or phase over time (Wacker and Witte, 2013).

### 2.7. Patients and Intra-Operative Measurements

For this study we use a test data set that consists of intra-operative measurements of eight patients with PD as shown in Table 1. At the time of operation the patients were withdrawn from their anti-parkinsonian medication for at least 12 h. Measurements were conducted during implantation of deep brain stimulation electrodes and LFP of up to five combined micro-and macroelectrodes were recorded with the INOMED ISIS MER-system (INOMED Corp., Teningen, Germany). The electrodes were arranged in Ben’s Gun-configuration (A: anterior, C: central, P: posterior, L: lateral, M: medial) with distances of 2 mm to the central electrode. The LFP analyzed in this paper were recorded with macroelectrodes of 1 kΩ impedance and sampled at 2456 Hz (Florin et al., 2013). Using microelectrode recordings we identified the locations, where single cell activity characteristic of the STN was observed (Benazzouz et al., 2002; Yang et al., 2014).

**Table 1:**
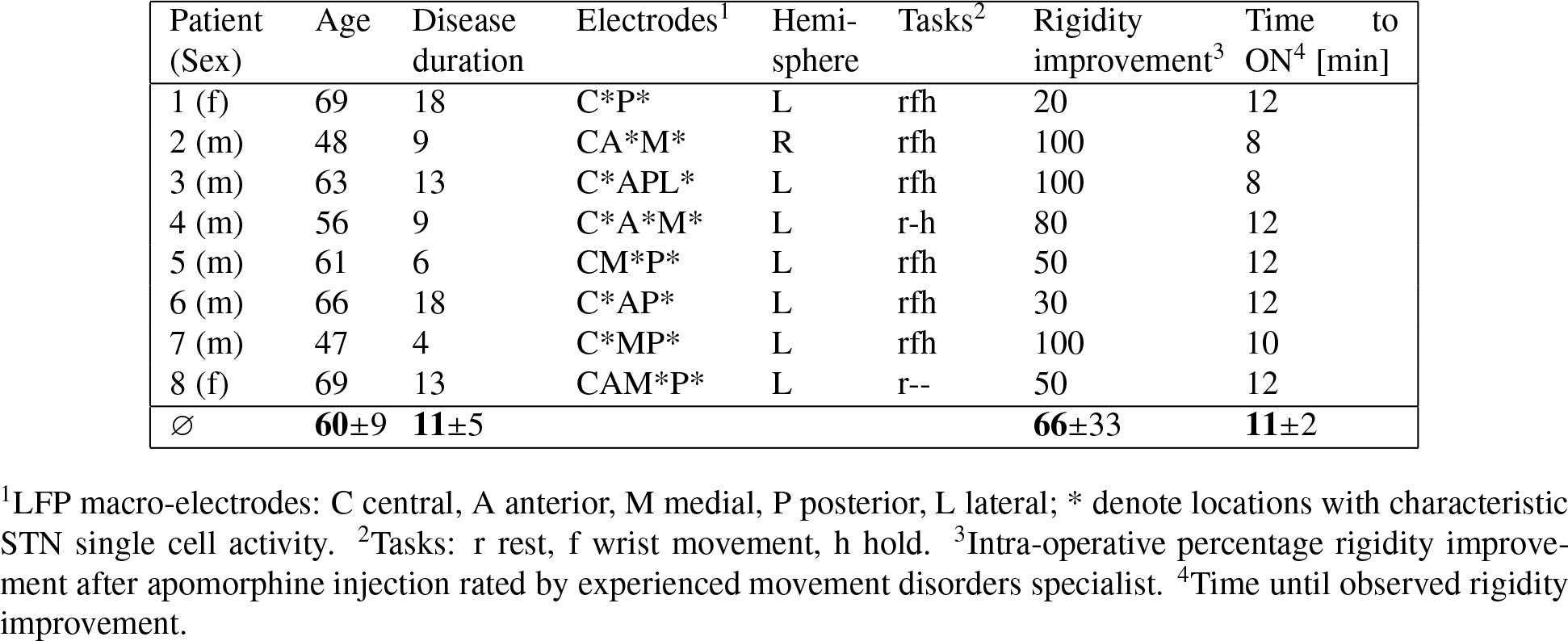
List of patients

The patients were measured before and after injection of apomorphine in order to assess the pathological activity in the LFP (see also Levy et al. (2001) for the effects of apomorphine). The patients were asked to perform simple motor tasks with the contralateral arm: a rest task, in which the patients relaxed their arm; an isometric hold task, in which the arm was held in a 45° angle; and a phasic wrist movement task, in which the patients were asked to form a relaxed fist and extend and flex their wrist at a pace of approximately 1 Hz. Each task was performed for 60 s. The complete set of rest/hold/wrist tasks was performed once before apomorphine injection (OFF) and was repeated after the medication effect had set in according to an experienced movement disorders specialist (ON). Every 2 min during the transition from OFF to ON the patients repeatedly performed the rest task. The ON state was identified from the improvement of the patients’ rigidity of the contralateral arm with respect to the baseline after electrode insertion (rated in 2 min intervals). Therefore, the improvement did not include the so-called "stun-effect", which can lead to an amelioration of symptoms possibly caused by microlesions from electrode insertion (Mann et al., 2009). Pre-operative the doses of apomorphine were individually adjusted according to clinical testing.

All patients gave written informed consent to the implantation of electrodes and the micro- and macroelectrode recordings. The study was approved by the local ethics committee (study no. 08-158) and conducted in accordance with the Declaration of Helsinki.

### 2.8. Relative and Absolute Changes of PSD

The relative power of each signal class *i* (incoherent, coherent and volume-conducted) was estimated according to 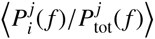 averaged in a specific frequency band of interest. The relative change of PSD from OFF to ON was defined as

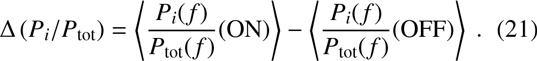

Note that for clarity we dropped the superscript *j* that denotes the electrode. We used this measure to compare signals before and after apomorphine injection because it accounts for changes of the baseline power. Such changes may arise because it takes several minutes to reach the ON state. However, the measure is prone to epiphenomenal changes as the sum of the relative differences over all signals *i* for a given electrode always amounts to zero.

In order to estimate movement-induced changes, we computed absolute differences according to

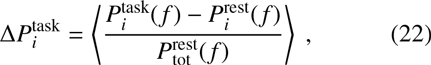

where 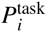 is the power of signal *i* during the specified task (hold, wrist) and 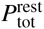 denotes the total power at rest. The task (hold, wrist) followed immediately after the rest measurement and, therefore, we did not expect strong changes of baseline power and hypothesized any changes of the signal to be caused by the movement task. We normalized these absolute differences to the total power at rest, so that 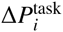 denotes the percentage change of signal class *i* induced by movement w.r.t. the total power at rest.

In our analyses of the electrophysiological response to apomorphine injection and movement we focused on the low and high beta bands (13–20 Hz and 20–30 Hz, respectively). We calculated the change of power for each electrode and estimated the significance of the results using a Wilcoxon signed rank test (due to nonnormal distribution) with a Bonferroni correction for multiple comparisons.

## 3. Results

We constructed synthetic time series with similar statistics as measured LFP and determined optimal parameters for the PCC, namely a suitable choice of *ω*_0_, *n*_*σ*_. We show that these parameters control the coherence and phase thresholds *C*_1%_ and Φ_*c*_, respectively. We then used the PCC to analyze beta band oscillations in LFP and estimate the LFP activity of local and remote electrophysiological activity within the STN area of patients with PD.

### 3.1. Comparison of Synthetic Time Series With Measured LFP

In a first step we compared our synthetic time series with measured LFP from within the STN of patients with PD during rest. The aim was to generate time series with known ground truth and similar first and second order statistics to measured LFP. An important parameter for that matter was the noise level controlled by *v*, which had to be set in relation to the power of the narrow band signals centered at *f*_*j*_. In real electrophysiological measurements this parameter is not known. Therefore, we compared the PSD of synthetic time series for different noise levels *v* with the PSD of real electrophysiological measurements in order to estimate a reasonable value for the noise level.

Figure 2 shows five seconds of a LFP measurement from a patient with PD within the STN (a) and a synthetic time series (b) according to Equation (19) with *v* = 3, i.e. the SD of the noise was three times larger than the amplitude of the sinusoidal signal. This corresponds to a signal-to-noise ration of −10 dB. It can be seen that the synthetic time series captured some characteristic dynamics of the LFP such as small steps (around 3.5 s) and intermittent spiking. In Figure 2c we compared the synthetic PSD with that of three PD patients. For a noise level of *v* = 3, we observed similar relative powers of narrow band oscillations with respect to the broad band background for both measured LFP and synthetic time series. Our synthetic time series with a noise level of *v* = 3 could thus reasonably well represent the first and second order statistics of measured LFP.

**Figure 2:**
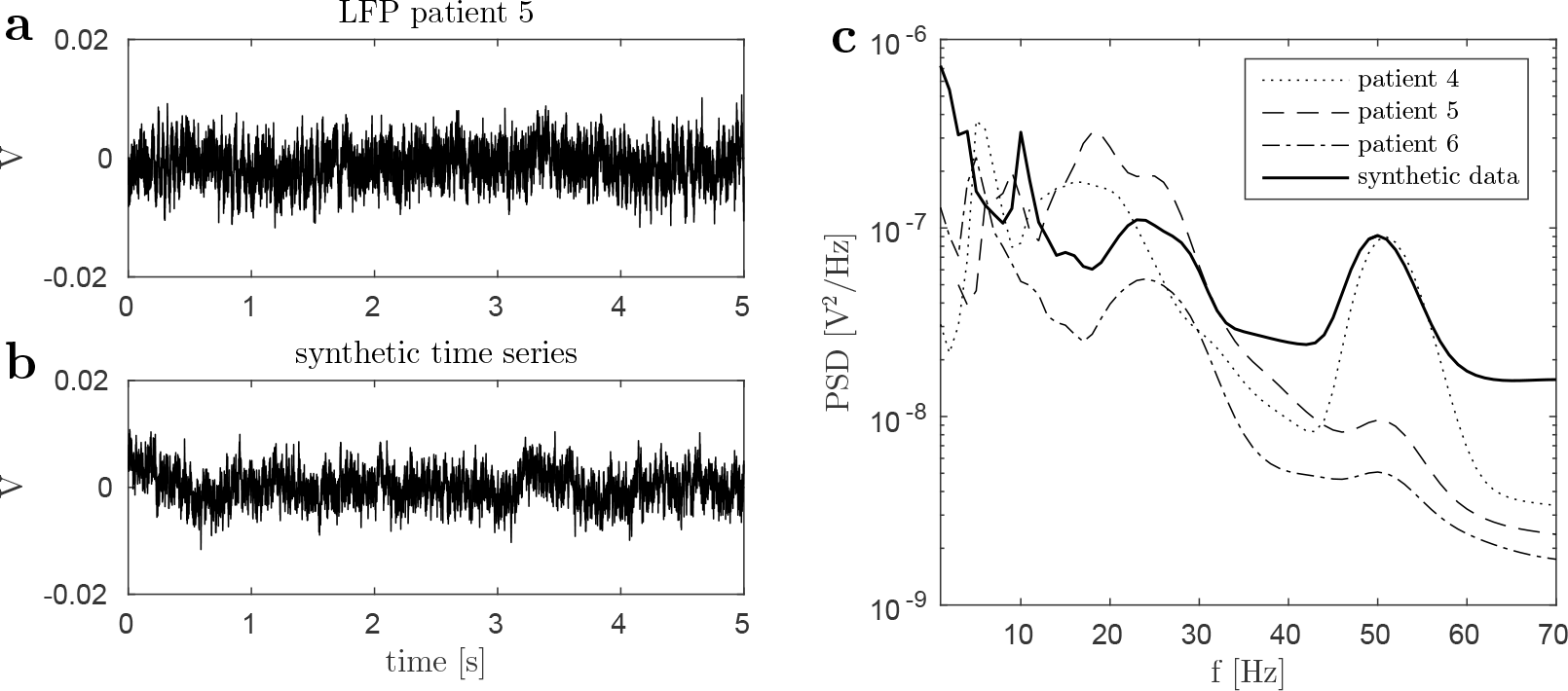
Time series of measured LFP of patient 5 within the STN (a) and synthetic time series (b) with a noise level of *v* = 3 for comparison. Amplitude and visible dynamics could be well replicated. b: PSD of LFP of three PD patients compared to the PSD of synthetic data showed similar relative power of narrow band peaks with respect to the broad band background.

### 3.2. Determination of Coherence Threshold

In order to determine a reasonable threshold for the time and frequency dependent coherence *C*(*f, t*) we constructed p.d.f. from pairs of white and pink noise. We tested the influence of averaging width *n*_*σ*_, wavelet parameter *ω*_0_ and frequency *f* on the estimation. In Figure 3a we show the p.d.f. of coherences *C*(*f, t*) obtained from 1000 pairs of pink noise at *f* = 10 Hz. It can be seen that the p.d.f. strongly depends on the averaging width. From visual inspection we found that the p.d.f. decreased sufficiently well for *n*_*σ*_ ≥ 4 in order to allow for a practical and reasonable determination of the significance threshold *C*_1%_. In Figure 3b we show the threshold *C*_1%_ as a function of *n*_*σ*_ for several wavelet parameters *ω*_0_. As expected, it decreased for increasing averaging widths and was independent of *ω*_0_ (Lachaux et al., 2002). The threshold was further independent of frequency and the type of noise (white or pink).

**Figure 3:**
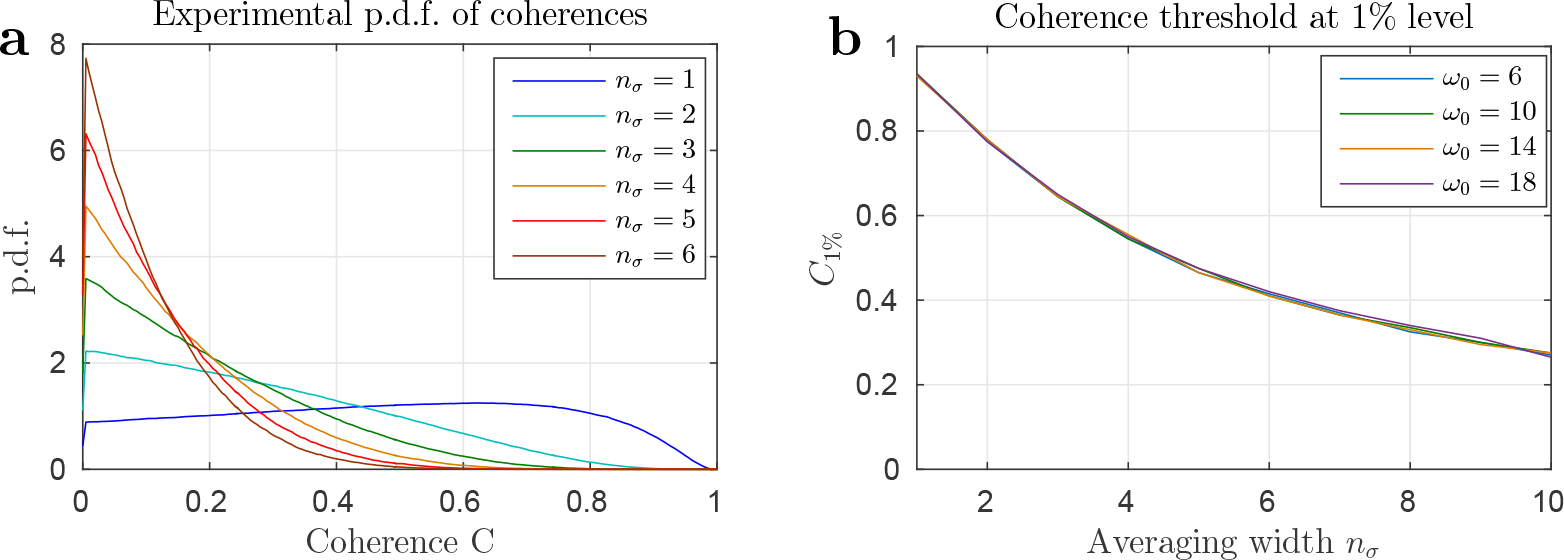
a: P.d.f. of coherences *C*(*f, t*) between 1000 pairs of pink noise at *f* = 10 Hz as a function of averaging width *n*_*σ*_. A sufficient decrease of the p.d.f. was observed for widths of *n*_*σ*_ ≥ 4. b: coherence threshold at a 1% level determined from the p.d.f., the threshold was independent of wavelet parameter *ω*_0_.

### 3.3. Determination of Phase Threshold

We estimated the phase resolution of the wavelet transform with non-phase-shifted synthetic time series generated by Equation (15). However, the observed phase differences between these two time series were expected to be distributed around the true phase difference of Φ = 0° due to the presence of noise and the finite resolution of the wavelet method. In Figure 4a we show the p.d.f. of the obtained phase differences Φ_*xy*_(*f, t*) at *f* = 20 Hz for several wavelet parameters *ω*_0_. Here we used a noise level of *v* = 3 and an averaging width of *n*_*σ*_ = 6. It can be seen that the p.d.f. are broader (phases less well resolved) for smaller wavelet parameters. Accordingly, the derived phase thresholds Φ_*c*_ were larger for smaller wavelet parameters.

**Figure 4:**
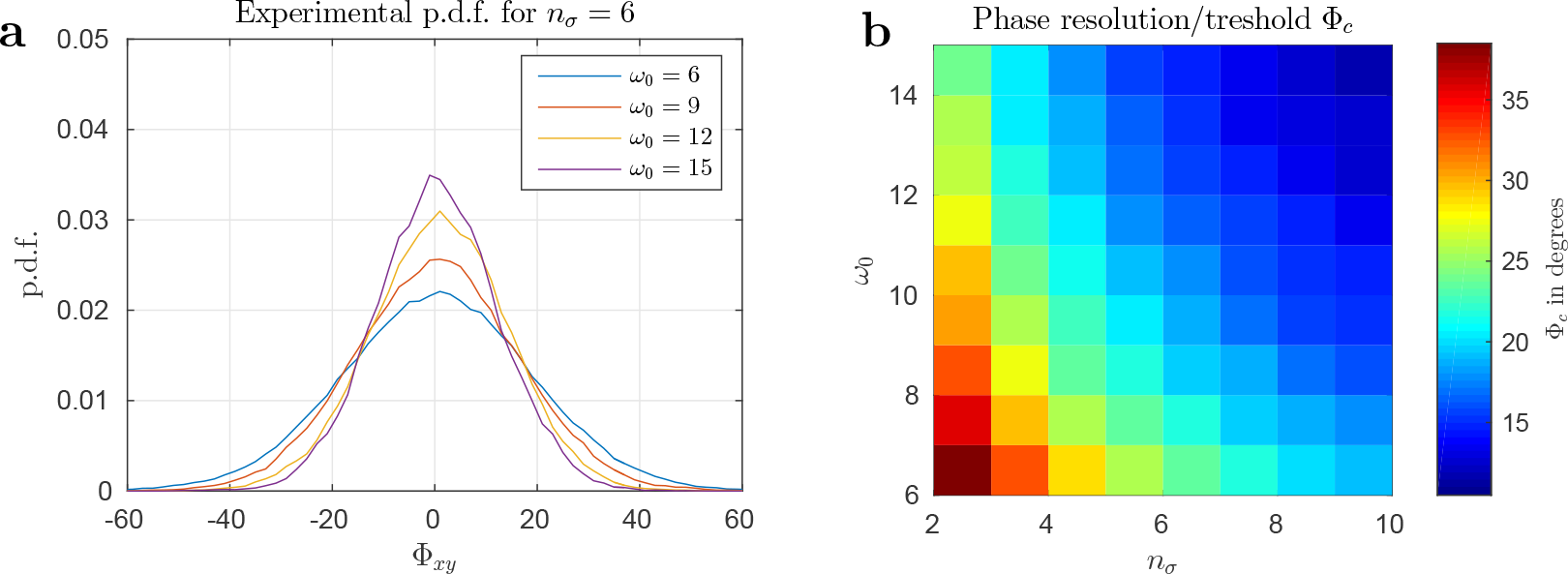
a: P.d.f. of observed phase differences Φ_*xy*_ between synthetic signals centered at *f* = 20 Hz in the presence of pink noise with *v* = 3 as a function of *ω*_0_. b: phase threshold as a function of *ω*_0_ and *n*_*σ*_. Generally, an increase of *ω*_0_ and *n*_*σ*_ resulted in better phase resolution (at the expense of poorer time resolution).

The averaging width *n*_*σ*_ had a similar effect on the phase threshold as the wavelet parameter. Namely, an increase of the averaging width resulted in a better phase resolution. This is shown in Figure 4b, where the phase threshold (color coded) is presented as a function of both wavelet parameter and averaging width *n*_*σ*_. For averaging widths of *n*_*σ*_ = 6 it resulted in Φ_*c*_ = 23.5° and Φ_*c*_ = 15.5° for *ω*_0_ = 6 and *ω*_0_ = 12, respectively. Note that Φ_*c*_ also depended on *n*_*σ*_ because we used time-averaged phases. The p.d.f. and thus the derived critical phase thresholds Φ_*c*_ were independent of frequency.

### 3.4. Choice of Optimal Parameters for PCC

The choice of optimal parameters crucially depends on the aim of the analysis. Here we focused on the analysis of LFP of patients with PD, where the beta band (13–30 Hz) plays an important role (Hammond et al., 2007). The beta band activity is highly dynamic and varies on scales of hundreds of milliseconds (Little et al., 2012). This poses a challenge on our method as the time resolution of the PCC, estimated by *σ* in Eq. (6), was generally larger than 200 ms. The time resolution as a function of *ω*_0_ and *n*_*σ*_ is shown in Figure 5 for the low beta band (*f* = 13 Hz).

**Figure 5:**
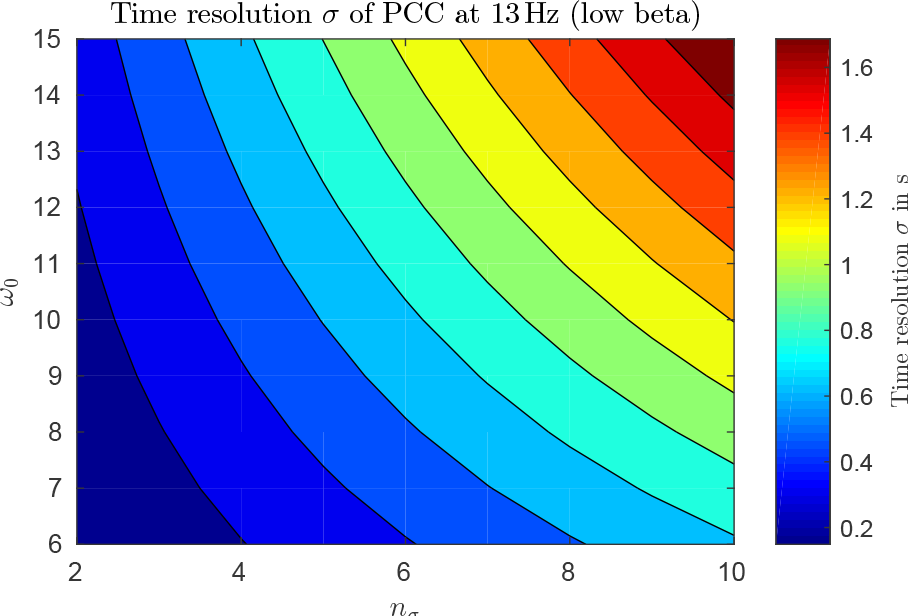
Time resolution *σ* as a function of *ω*_0_ and *n*_*σ*_ for low beta band frequency *f* = 13 Hz. Blueish colors denote resolution of less than 1 s.

In order to determine optimal parameters we looked for a balanced choice between time and phase resolution (Figures 4b and 5, respectively). We found that *ω*_0_ = 12 and *n*_*σ*_ = 6 was a reasonable choice for the further analyses in the beta band as it allowed a high phase resolution (Φ_*c*_ = 15.5°) with a temporal resolution of *σ* = 0.37–0.88 s. Also, the wavelet parameter was large enough to guarantee a high frequency resolution (Δ*f* / *f*(*ω*_0_ = 12) < 1.14), which was important to discriminate between the low (13–20 Hz) and high (20–30 Hz) beta band. The coherence estimate for this choice of parameters resulted from a temporal averaging of more than 2.4 ⋅ *n*_*σ*_ *sf* ≈ 27 periods of the frequency under consideration with a threshold of *C*_1%_ = 0.41.

### 3.5. Accuracy of the PCC

In order to test how much power the PCC assigned to each signal class (incoherent, coherent and volume-conducted) we used 500 pairs of synthetic time series according to Equation (15) with a signal centered at 20 Hz and applied variable phase shifts 0° ≤ Φ_12_ ≤ 40° between the two time series. The result is shown in Figure 6, where we plot the relative power of the coherent (*P*_coh_/*P*_tot_) and volume-conducted signal (*P*_vc_/*P*_tot_). For no phase shift our method assigned 77 ± 14% of the total power to the volume-conducted signal. For a phase shift of Φ = 25° we found that 74 ± 17% of the signal was assigned correctly to the coherent signal increasing to 94 ± 7% for a phase shift of 40°. For large phase shifts the local coherent signal could thus be identified more accurately than the volume-conducted signal. On average only 2% of the power was incorrectly assigned to the incoherent class.

**Figure 6:**
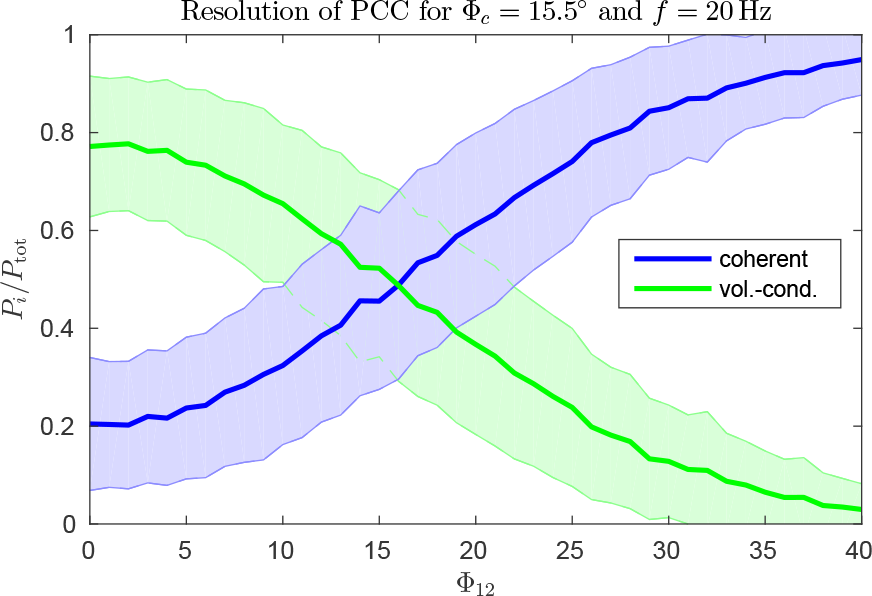
Resolution analysis of PCC for varying phase shifts ΔΦ with a fixed phase threshold of ΔΦ_*c*_ = 15.5°. The larger the phase shift the more power was assigned to the local coherent signal. At zero phase shift there was still about 20% power in the local coherent signal due to noise. Colored areas indicate SD from 500 trials.

As a proof of concept we applied the PCC to non-stationary time series with temporal variations of amplitude, frequency and phase. For that matter we used composite synthetic time series as defined in Section 2.6. Figure 7a shows the classification of the wavelet coefficients in time-frequency space for a single run. The signal classes are color coded and the black line denotes the cone of influence. It can be seen that PCC correctly identified coherent signals at 10 Hz, 20–30 Hz and 50 Hz before an incoherent background. The phases of the coherent signals could be well resolved so that the PCC correctly distinguished between local coherent and volume-conducted signals. Only very few coefficients were falsely classified.

**Figure 7:**
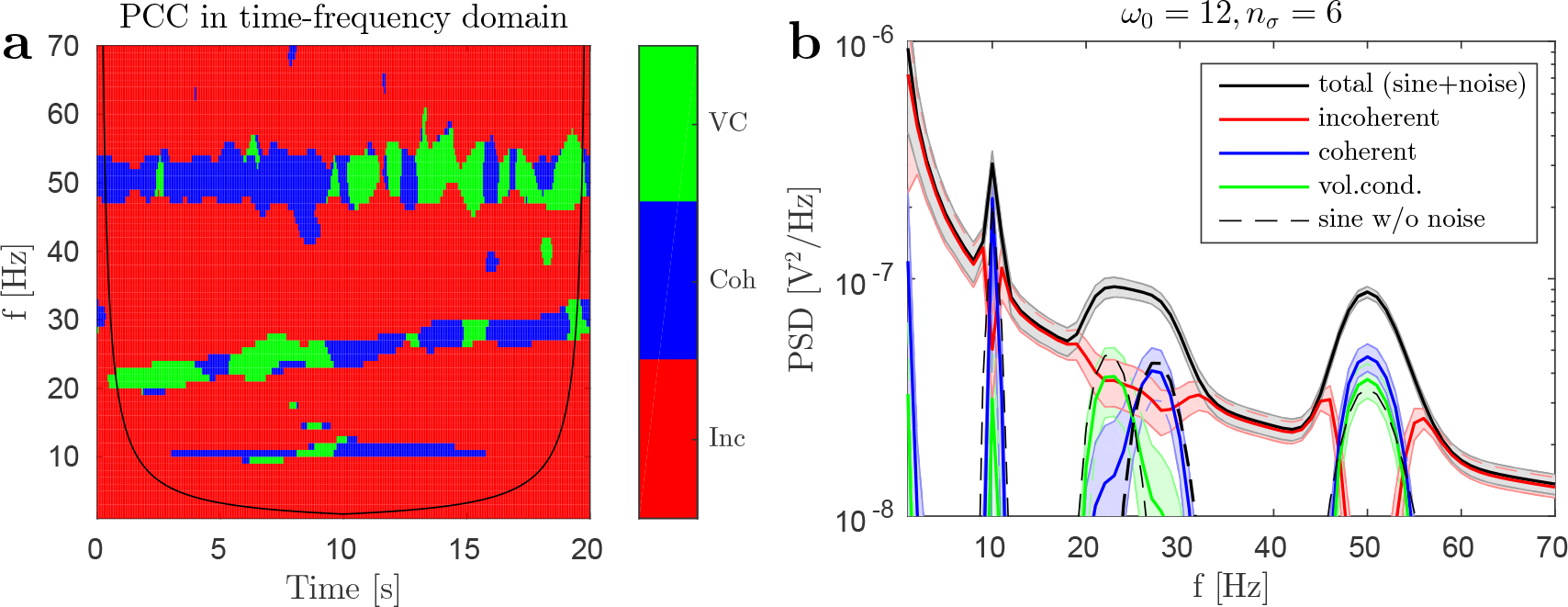
PCC for synthetic time series with signals of varying amplitude, frequency and phase. a: result of a single trial in time-frequency domain, b: ensemble averaged PSD (mean±SD). PCC could well classify the signals at 10 Hz, 25–30 Hz for *t* > 10 s and 50 Hz for *t* ≤ 10 s as local coherent, and the signals at 20–25 Hz for *t* ≤ 10 s and 50 Hz for *t* > 10 s as volume-conducted. The dashed black line in the right panel denotes the true signal power, which PCC generally reproduced within error bars.

The corresponding PSD averaged over 1000 trials are shown in Figure 7b. The solid lines denote the means and the colored areas denote the SD of the ensemble. The black dashed lines denote the power of the signals without additional noise and therefore reflect the best estimate. The PCC reproduced the best estimate generally within error bars except for the coherent power at 50 Hz, which was slightly overestimated. The SD were small compared to the mean with on average SD/mean = 20% for the data visible in the right panel of Figure 7 (the ratio naturally increases when lower powers are also considered because the mean of PCC signals approaches zero for some frequencies).

### 3.6. Case Study: Detection of Local Tremor Clusters

Here we apply the PCC to the LFP of patient 3 during a tremor episode at rest and off medication. The incoherent, coherent and volume-conducted power of the LFP were averaged according to Equation (13). In Figure 8a we show results of a case study for channel A (w.r.t. reference electrode P). In order to show the temporal dynamics of the tremor we present the LFP time series in gray and the power at 5 Hz in black. It can be seen that the tremor power increased around *t* = 10 s after measurement onset. From this time on - as shown in Figure 8b - the PCC classified the tremor throughout as local coherent. The power of the local coherent signal is visible in the PSD in Figure 8c. It was the dominant signal in the tremor peak at 5 Hz, which confirmed the presence of multiple local tremor clusters. Interestingly the power at double tremor frequency was mostly incoherent. This may indicate differing wave forms at the two measurement sites as the waveform is determined by higher harmonics in frequency space.

**Figure 8:**
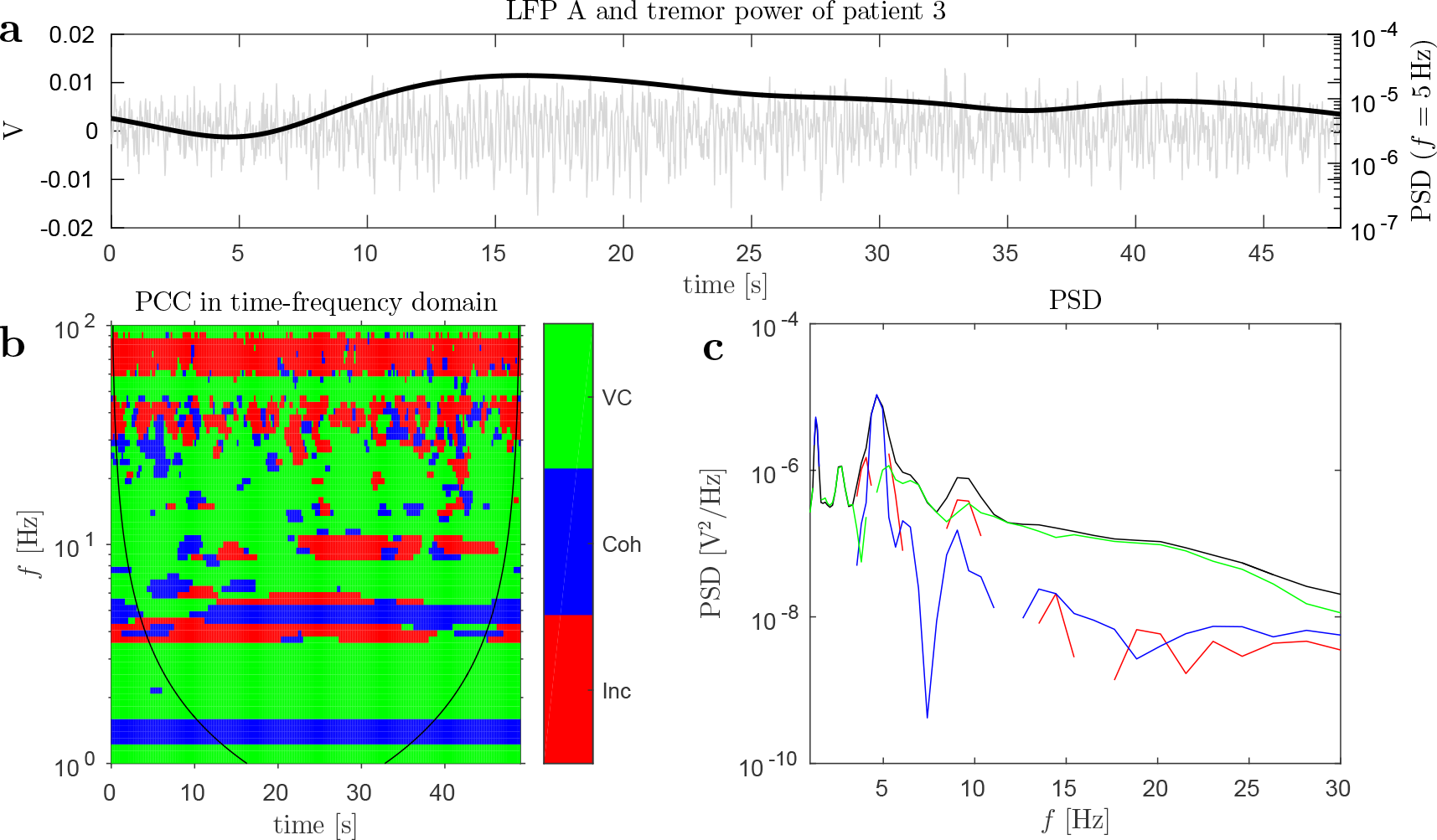
Case study of patient 3 during tremor episode at rest and off medication. a: LFP A (gray) and PSD at tremor frequency (*f* = 5 Hz, black). b: PCC in time-frequency domain with color coded signal classes. c: Time-averaged PSD of incoherent, coherent and volume-conducted power. The tremor started at around *t* = 10 s at 5 Hz and consisted of coherent power, which indicates multiple sites of local activity.

### 3.7. Group Analysis: Relative Power of Signal Classes

We analyzed the power spectral densities of LFP with characteristic STN activity for the patients presented in Table 1. On average the intervals suitable for analysis after artifact removal were of length 42 ± 19 s, 49 ± 11 s and 49 ± 16 s, for rest, hold and wrist, respectively. Due to incorrect performance of the active task, we had to exclude some trials from the analysis, which resulted in data sets of 8 patients for rest, 7 patients for hold and 6 patients for the wrist movement task. For each OFF and ON measurement this led to a total number of 17, 15 and 12 monopolar recordings for rest, hold and wrist movement, respectively.

In a first step to analyze the signal classes obtained from PCC we calculated their relative contribution to the total power. In Figure 9 we show the mean relative power during rest and off medication with corresponding standard error of the mean (SEM) denoted by shaded areas. The other tasks (hold, wrist) resulted in similar distributions of power. The relative contribution of the incoherent signal increased strongly from frequencies *f* < 40 Hz, where it contributed 10–30% of the total power, to frequencies *f* > 70 Hz, where it contributed up to 90%. The local coherent power was largest at frequencies below 15 Hz, where it contributed about 30% of the power, and remained at approximately 20% in the beta and low gamma band (*f* ≤ 40 Hz). At frequencies *f* > 60 Hz it contributed less than 10% to the total signal. The volume-conducted signal was generally the largest fraction for frequencies below 60 Hz and peaked in the beta band (13–30 Hz), where it contributed about 70% to the total signal. For frequencies *f* > 60 Hz its contribution dropped to less than 10%. At even higher frequencies *f* > 100 Hz the percentage contributions stayed approximately the same as those observed at 80 Hz with 90% incoherent and 5% coherent and volume-conducted power each. Note that the relative power at 50 Hz was distorted by power line noise reflected in an increased volume-conducted signal and decreased incoherent power.

**Figure 9:**
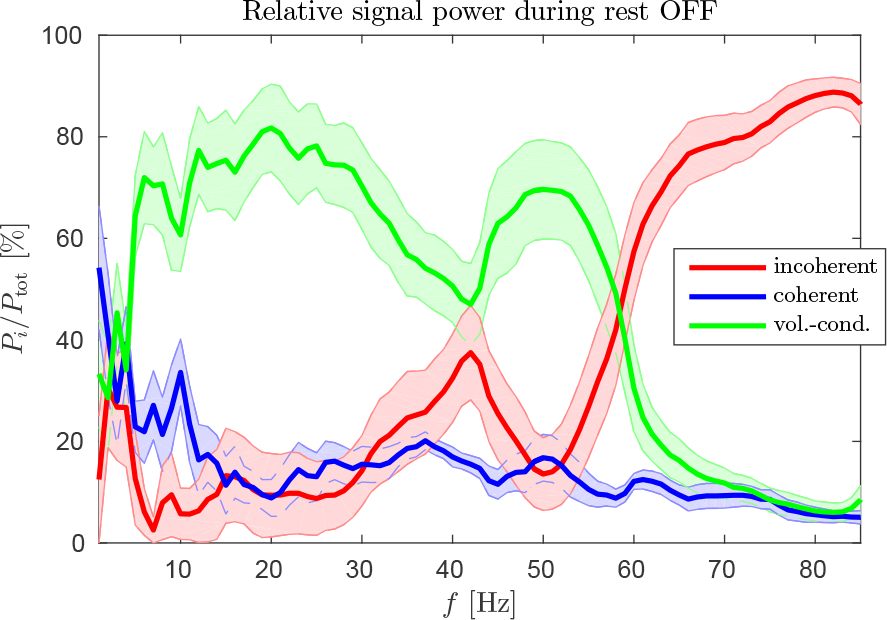
Relative PSD of local incoherent, coherent and volume-conducted signals during rest off medication for eight PD patients (mean ± SEM). The incoherent signal increases with higher frequency (*f* > 60 Hz) reflecting more local activity. The volume-conducted signal is largest up to about 40 Hz. The peak at 50 Hz is caused by line noise.

### 3.8. Electrophysiological Response to Apomorphine Injection

The relative change of spectral power during rest induced by apomorphine injection was analyzed for all locations with characteristic STN single cell activity. In the left and right panel of Figure 10 we show the relative changes according to Equation (21) for the low and high beta band, respectively. Asterisks denote 5% significance before (*) and after (**) Bonferroni correction for six comparisons. We estimated the significance with a nonparametric Wilcoxon signed rank test. PCC signals showed that the share of incoherent power to the total signal significantly decreased from OFF to ON (*p* = 0.0034, test statistic *W* = 4, number of samples *N* = 17) in the low beta band (although not very strongly for many electrodes) while the volume-conducted part significantly (*p* = 0.031, *W* = 31, *N* = 17) increased. Opposing trends were observed in the high beta band although not significant.

**Figure 10:**
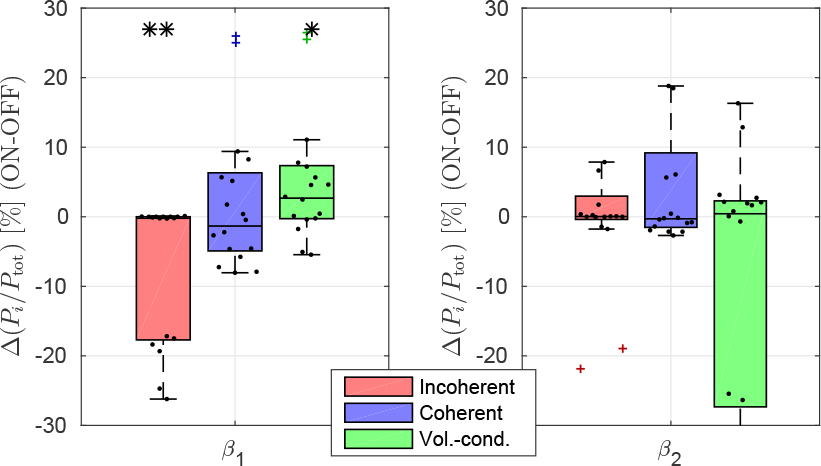
Relative change of spectral power Δ (*P*_*i*_/*P*_tot_) (dots and crosses) from OFF to ON for all electrodes with STN activity averaged in low beta (*β*_1_ = 13–20 Hz, left) and high beta (*β*_2_ = 20–30 Hz, right). Asterisks (*) and (**) denote 5% significance before and after Bonferroni-correction for six comparisons, respectively.

### 3.9. Changes Induced by Motor Tasks

The absolute changes in high beta band induced by a holding task and wrist movement are shown in the top and bottom panel of Figure 11, respectively. The local coherent signal changed significantly in power on medication. It was responsible for an average increase of 3.1% of total power induced by the hold task (*p* = 0.022, *W* = 20, *N* = 15) and an average decrease of 5.3% of total power induced by wrist movement (*p* = 0.0034, *W* = 4, *N* = 12), where the latter remained significant after Bonferroni correction for twelve comparisons (three classes OFF and ON in low and high beta band). The same analysis of changes in the low beta band yielded no significant results. Using relative changes of power instead of absolute changes led to similar results: the coherent signal increased about 1.7% during the hold task although not significantly (*p* = 0.14, *W* = 33, *N* = 15) and decreased significantly (before Bonferroni correction) about 3.9% during wrist movement (*p* = 0.027, *W* = 11, *N* = 12).

**Figure 11:**
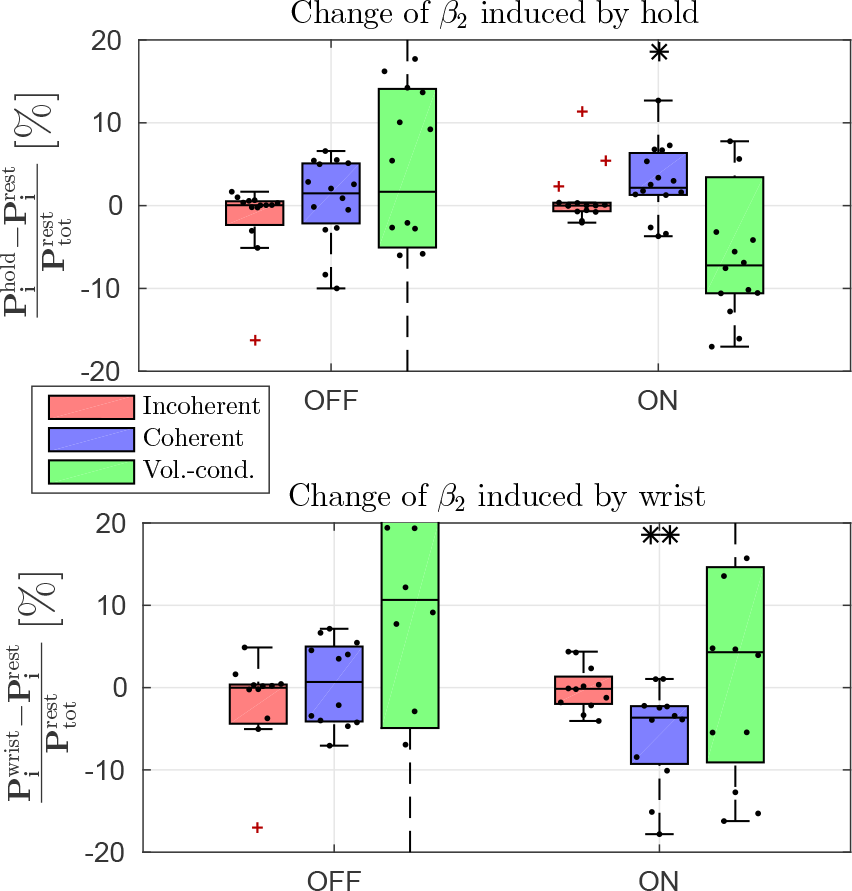
Absolute changes induced by movement (top panel: hold, bottom panel: wrist movement) in the high beta band (20–30Hz) with respect to rest off and on medication (dots and crosses) normalized by total power at rest. Asterisks (*) and (**) denote 5% significance before and after Bonferroni-correction for twelve comparisons, respectively.

## 4. Discussion

We presented the phase-coherence classification, a novel wavelet-based method to separate LFP into local incoherent, local coherent and volume-conducted signals based on the pairwise phase and coherence characteristics. We applied this new technique to synthetic time series and LFP from patients with PD. We showed that the PCC could well capture non-stationary features and that the signal classes for PD patients were differently modulated by medication and movement: incoherent/volume-conducted signals significantly decreased/increased in the low beta band after apomorphine injection, while coherent signals showed significant movement-induced changes on medication. These results suggest that the signal classes reflect physiologically and functionally different neuronal network activity. The classification of (in)coherent and volume-conducted signals may thus provide useful information to distinguish between movement-induced and medication-induced (OFF/ON) LFP changes.

### 4.1. Methodological Considerations

The synthetic time series used in this study were constructed to reflect the statistical characteristics of LFP measurements. This has been done phenomenologically by visual comparison of time series and power spectra. Higher order statistics of measured LFP such as skewness and kurtosis were found to be highly variable between measurements. However, on average these statistics were similar to those of synthetic time series. In order to imitate observed narrow band peaks in the PSD of measured LFP we used sinusoidal signals with time-varying amplitude, frequency and phase.

We carefully tested the phase-resolution of the wavelet transform in order to determine a suitable phase threshold Φ_*c*_ for the PCC. Note that the PCC is not able to discriminate between remote electrophysiological activity (volume-conducted) and local activity with a phase difference smaller than this threshold. This type II error is inherent to the PCC and was considered in the interpretation of our results (see Section 4.2). However, compared to the imaginary part of the coherency (Nolte et al., 2004), the PCC explicitly considers phase information and can thus resolve even minimally significant coherent signals. The threshold was defined so that 80% of the observed phases from a non-phase-shifted signal are correctly classified. The threshold was shown to depend on the wavelet parameter *ω*_0_ and averaging width *n*_*σ*_, which need to be chosen according to the individual scientific question at hand. The choice also depends on the signal-to-noise ratio of the data. In the presence of strong noise *n*_*σ*_ needs to be larger in order to estimate the coherence reasonably well (Lachaux et al., 2002).

We tested how well the PCC performed for non-stationary data using composite signals of varying amplitude, frequency and phase. We quantified how much power was assigned to each class and showed that PCC could well reproduce the true signals even in the presence of noise. This is of great importance in the analysis of electrophysiological data and makes the PCC suitable to detect non-stationary phenomena (see Gross et al., 2000; Bokil et al., 2007; Bigot et al., 2011; Wacker and Witte, 2013, for other methods that take non-stationarity into account).

The PCC assigns exactly one class to every coefficient in time-frequency space. This means that the PCC only classifies the signal that is strongest at that time and frequency while weaker signals may remain undetected in the background. An example of such behavior was presented in the right panel of Figure 7. Here the incoherent signal, which was present at all times and frequencies, decreased to zero at 50 Hz because the coherent and volume-conducted signals were stronger. This winner-takes-it-all characteristic is an important limitation of the PCC. However, the influence of this characteristic on non-stationary signals can be decreased by a high time resolution.

The main advantage of the PCC over conventional monopolar or bipolar recordings is the separate analysis of local incoherent, local coherent and volume-conducted signals, which may represent functionally separated network activity. The different signal classes can further be associated with possible locations of electrophysiological activity. The possibility to analyze the volume-conducted signal separately may give rise to further insights about remote activity or local activity with nearly no phase difference. The PCC thus allows for a better spatial and functional interpretation of monopolar signals.

Finally, we note that the PCC scheme should hold for any signal generated from the integrated activity of a large population of neurons, i.e. not only LFP but also EEG and MEG, as long as the quasi-static approximation holds. In EEG and MEG analyses the PCC may be used in order to minimize spatial leakage caused by a limited spatial resolution. Compared to other measures (Nolte et al., 2004; Hipp et al., 2012; Colclough et al., 2015) the PCC preserves the volume-conducted signal and explicitly takes the phase information into account. Further it can be adjusted to the problem at hand through the parameters *ω*_0_ and *n*_*σ*_. A leakage free functional connectivity measure, e.g., can be estimated by averaging coherences associated with phases Φ > Φ_*c*_ to eliminate volume-conduction effects.

### 4.2. Clinical and Neurophysiological Discussion

In a case study of a patient at rest during a tremor episode we showed that the tremor signal at 5 Hz consisted of local coherent power. This confirmed the existence of local tremor clusters in the STN area as has been estimated from coherence analysis by Reck et al. (2010). Interestingly, the activity at double tremor frequency was mostly incoherent. The fact that we observed incoherent power at double frequency for many other electrode combinations and their respective reverse combinations (X→Y and Y→X) may be explained by different waveforms at single tremor frequency at both electrodes. This would lead to different harmonics and thus to non-significant coherence at double tremor frequency.

We chose the parameters in our study of the test data set in order to allow for a detailed analysis of the low and high beta band activity. In the frequency range of 13–30Hz this led to a temporal resolution of *σ* = 0.38–0.88 s. This was somewhat larger than the typical duration of beta bursts, which are on the order of hundreds of milliseconds (Courtemanche et al., 2003; Androulidakis et al., 2006; Little and Brown, 2012). Thus, our coherence estimates included the variability of instantaneous beta band activity caused by these bursts.

Using PCC we were able to estimate the average relative power of the signal classes in the patient cohort. The beta band activity within the STN consisted mostly (~90%) of coherent signals generated by local and possible remote activity. Signals at high frequencies (*f*>40 Hz) mainly showed local incoherent activity. It is well known that the spatial reach of LFP is frequency-dependent (Stinstra and Peters, 1998; Lindén et al., 2010; Buzsáki et al., 2012; Leski et al., 2013), however, this is to our knowledge the first time that the amount of local and remote activity within the STN was quantified as a function of frequency.

The decrease of low beta band power from OFF to ON was partly caused by local incoherent activity that showed a significant decrease during rest. At the same time the volume-conducted signal increased significantly, however, only before Bonferroni correction. This might indicate that the local activity within the STN decreased thus rendering the nucleus less opaque to remote activity. However, in the high beta band this was not the case and volume-conducted signals tended to decrease rather than increase. Unfortunately, it remained unclear where exactly the activity of these non-phase-shifted (volume-conducted) signals stems from. According to our discussion of the type II error in Section 4.1, there are two possible interpretations of the volume-conducted signal: 1) it reflects remote activity external to the STN or 2) it reflects local in-phase activity with phase differences of less than 15.5° between electrodes. In case of pathological synchronization within the STN (Brown, 2007; Hammond et al., 2007; Kuhn et al., 2009) latter interpretation may be correct, i.e. large parts of the STN encompassing several electrodes oscillated in-phase.

For the local coherent signal movement-induced changes in the high beta band for both hold and wrist movement were in agreement with previous observations made from bipolar measurements, namely a decrease during phasic movement (Cassidy et al., 2002; Levy et al., 2002; Kuhn et al., 2004; Alegre et al., 2005) and an increase during isometric contraction (Baker et al., 1997; Jenkinson and Brown, 2011). The movement-induced changes of the local coherent signal were only observed on medication and may thus reflect healthy beta band modulation in response to motor function. This indicates that coherent signals might be generated by a functionally separated network that could only be (de)synchronized in the physiological state.

In tests with bipolar recordings that we conducted for comparison (not shown) the movement-induced change of power in the high beta band induced by wrist movement was slightly larger off medication than on medication although not significant. This is generally in agreement with observations by Cassidy et al. (2002) and Priori et al. (2002) who found stronger beta band desynchronization off medication. However, in trial based analyses Doyle et al. (2005) observed larger event-related desynchronization (ERD) of beta band activity on medication, while Alegre et al. (2005) observed the same ERD off and on medication. These contradicting observations might be explained by the different modulation of incoherent and coherent power: during wrist movement the incoherent signal decreased more strongly off medication (Δ*P* = −2.7% OFF vs. Δ*P* = 0.5% ON) while the local coherent signal decreased more strongly on medication (Δ*P* = 1.0% OFF vs. Δ*P* = −5.3% ON). As the bipolar recording is not able to discriminate between those two signals, this may lead to varying results depending on the baseline power of each signal class.

Our results further supported previous observations that the low beta band is more susceptible to medication-induced changes while the high beta band better reflects movement-induced changes (Priori et al., 2004; Foffani et al., 2005). The PCC added useful information to this phenomenon by attributing the respective changes to different signal classes, namely the incoherent power to medication-induced and local coherent power to movement-induced changes.

## Acknowledgements

This investigation was supported by the Competence Area 3 of the University of Cologne as part of the Excellence Initiative and the Clinical Research Group 219 (KFO219) of the German Research Foundation (DFG). The authors would like to thank Benjamin Mille and Tobias Koch for their help in visual inspection and preprocessing of the data.

